# Use of Artificial Intelligence for Automated Detection and Surveillance of Red Imported Fire Ants Nests

**DOI:** 10.1101/2023.05.26.542461

**Authors:** Xin Su, Guijie Shi, Jiamei Zhong, Yuling Li, Wennan Dai, Guohua Xu, Eduardo G. P. Fox, Hualong Qiu, Zheng Yan

## Abstract

The red imported fire ant, *Solenopsis invicta* Buren is a destructive invasive species that has spread around the world. Early detection of *S. invicta* nests is critical for effective monitoring and control in invaded regions. This study presents a novel surveillance system for *S. invicta* nests combining artificial intelligence and robotic dogs. The system was designed with intelligent recognition algorithms to accurately identify *S. invicta* nests. With a precision rate of 95%, the system yielded efficient detection of *S. invicta* nests with increased sensitivity and low missing and false discovery rates, which can aid or even replace humans in locating and delivering pesticides to fire ant nests in open fields to effective control of this invasive pest.

## 1 Introduction

Invasive non-native species pose a significant threat to global biodiversity, public safety, and economic stability^[1]^. The Red Imported Fire Ant (*Solenopsis invicta*), or RIFA for short, was accidentally introduced to the United States in the 1930s and has since caused extensive environmental damage and economic losses. Its spread across the American South and into northeastern Mexico results in billions of dollars in damages each year due to property destruction, pest control, and containment efforts^[2]^.Unfortunately, RIFA populations have been introduced through international shipping into several countries, including Australia (2001), China (2004), Japan (2017), and South Korea (2018)^[3]^. Standard approaches to controlling or containing these populations typically involve the use of pesticides^[4]^, but these can have harmful effects on local ecosystems.As a result, it is essential to establish clear guidelines surrounding pesticide usage, such as dosage, timing, and application windows, to reduce negative side effects and target non-native species more effectively.

Published research on RIFA has primarily focused on its physiological characteristics, invasion risk, and integrated pest management strategies^[5–8]^. Integrated pest management relies heavily on monitoring invasive species populations to deliver effective control measures in a timely manner. One widely-used method for estimating RIFA density involves tallying the number of active nests per unit of area^[9]^. However, identifying RIFA nests visually can be challenging due to their diverse and non-distinctive physical profiles, and confirmation of RIFA presence often requires molecular techniques^[10]^.Various attempts have been made to develop technologies for automatically detecting RIFA nests, including through the use of aerial imagery^[10,11]^, drones^[6]^, and trained dogs^[12,13]^. Nevertheless, the scalability of these methods is often hindered by the need for specialized machinery and expertise.

To supplement existing in-field surveillance efforts, using sensing robots has been proposed as a feasible strategy^[2]^. This approach heavily relies on real-time performance algorithms to detect RIFA nests in the open field. Artificial intelligence (AI) technology, with its ability to automate target detection and searching, has experienced growing applications in this regard, surpassing traditional image-processing methods^[14,15]^. Among current algorithms like R-CNN, YOLO, and SSD^[16–18]^, each has its advantages and disadvantages. While R-CNN offers high accuracy, it lacks the ability for rapid detection in real-time situations. Alternatively, while SSD can distinguish between large-sized targets with high accuracy, its performance drops when recognizing smaller size targets.Comparatively, YOLO can classify objects quickly and accurately by regressing image information, classifying objects, and labeling themes with sufficient precision. As the acronym suggests (‘You Only Look Once’), YOLO uses a single CNN network to detect and classify an object without inspecting the entire subject or the surrounding area ^[17]^.In 2021, Kuang-Shi Technology (MEGVII) upgraded YOLO algorithms with an innovative decoupled head, mosaic and MixUp augmentation strategies, anchor-free manner, and improved small size target recognition accuracy, introducing YOLOX^[19]^. The YOLOX model won 1^st^ place for the Streaming Perception Challenge (Workshop on Autonomous Driving at CVPR 2021) processing at 68.9 frames per second (FPS) on Tesla V100^[19]^, indicating reliable image processing capacity for practical applications.

Although still in their early stages of development, robotic dogs have enormous potential to revolutionize how people interact with their environment and each other. Typically, robotic dogs incorporate various sensors such as cameras, gyroscopes, accelerometers, sonar sensors, and force sensors enabling them to carry out tasks like fetching objects and opening doors while interacting with their surroundings^[20]^. Additionally, they possess a level of autonomy that allows them to navigate through complex environments without relying on human input^[21]^. Furthermore, they may be equipped to detect and mirror human emotions, providing an alternative source of companionship and comfort for their owners^[22]^.In the field of pest management, four-legged robotic dogs offer a promising solution to suppressing the Red Imported Fire Ant (RIFA) population despite various obstacles and challenges such as unreliable power sources or unstable robot navigation across bumpy, soft, and cliff terrain. When compared to wheeled robots, legged systems offer better performance and mobility across irregular grounds and, with improved image recognition and AI technology, should have the capability to effectively recognize and distinguish RIFA nests^[23]^.The application of robotic dogs is gaining increasing popularity and development, presenting their great potential for numerous industries such as medical assistance, environmental exploration, companionship, pest management, and more^[24]^.

To achieve this, we utilized the CyberDog robotic platform manufactured by Xiaomi Global. This open-source robot is equipped with multiple cameras, image sensors, and a powerful Nvidia Jetson Xavier AI processing unit with a data storage capacity of 128 GB^[19]^. Our approach combined the CyberDog robotic platform with YOLOX technology to develop a pilot program for automatic sensing detection of RIFA nests. This program aims to improve pest population monitoring and enable more efficient timing for pesticide control.

The Red Imported Fire Ant (RIFA) populations are rapidly spreading into previously uninfected areas in China^[25]^. Automatic sensing could provide invaluable assistance in monitoring fire ant population density, especially in locations that are difficult to access for pest control personnel such as airport runways. The objective of our project was to assess the feasibility of using commercially-available robots along with open-source software to automatically detect and register RIFA nests in open fields in China. Our results indicate that current state-of-the-art AI technology can detect RIFA nests with a success rate of over 90%, making it possible to take proactive measures in order to stop the invasion. The main sources of detection error were related to smaller nests that have been overlooked and situations where mated RIFA queens were still developing their nests.

## 2 Materials and methods

### 2.1 Image Data collection

We constructed a database of RIFA fire ant nest images from China, Brazil, and the United States. The images were collected from various sources, including iPhone (models 6 and 13) cell phone cameras, footage stills from a GoPro HERO 95K Ultra HD, and 404 internet-sourced images obtained using a web crawler. We utilized both paid and free stock photos to build our dataset. For training and implementing the YOLOX model, we used three sets of images (Table 1): (a) 1,519 raw digital images of RIFA nests taken under different weather conditions and backgrounds; (b) after annotation, the images were divided into a training set (n=1118) and a test set (n=401); (c) we also included 100 images of nests from other ants as negative controls. All images were saved in JPG format. By utilizing images from multiple sources and locations, we aimed to create a diverse and comprehensive dataset that would improve the accuracy and robustness of our object detection model.

**Table 1.** Images of ants and their nests obtained for YOLOX model training

| Image Type | *Solenopsis* fire ants | Other ants |
| --- | --- | --- |
| Ant nests from China | 1519 | 10 |
| Ant nests from US and Brazil | 47 | 100 |
| Individual ant images | 22 | 5 |

### 2.2 Data pre-processing and annotation

To accurately identify the Red Imported Fire Ant (RIFA) nests in our images, we utilized the annotation tool LabelImg^[26]^. Each nest was marked with a rectangular shape that precisely indicated its location and size by touching the borders of the nest, resulting in clear measurements for both width and height. We carefully labeled each object before saving the individual annotations in the COCO format, which is compatible with the YOLOX model. This method ensured that our data was properly annotated and prepared for further analysis.

### 2.3 the YOLOX model Training

To train YOLOX models on a local computer with a GEFORCE RTX 3060 GPU, we used a dataset and a command script provided in the YOLOX directory. Pre-trained COCO weights were utilized in the training process, which lasted for 400 epochs and included a 5-epoch warm-up. We used stochastic gradient descent (SGD) as well as a weight decay of 0.0005. During warm-up, the learning rate was set to 0, while the basic learning rate was set to 0.00015625. The ratio of the final learning rate was set at 0.05. To measure FPS (Frames Per Second) and latency, we used FP16-precision and batch=1. We outputted training results per interval of 30, and validated per epoch = 1.After training, we tested the model on the website http://391x54r871.wicp.vip. Overall, we believe our approach represents an effective and efficient way to train and test YOLOX models for object detection.

### 2.4 Setting up of the model, web page detection and real-time identification of ant nests

We set up the YOLOX program by uploading the model, compressed as a file, to the CyberDog server. We then installed the corresponding Python environment and CUDA configuration on the server. The robot dog’s camera was connected to the program using OpenCV to capture real-time images, which were processed in the program for the real-time detection and identification of ant nests. To make the model accessible to multiple users, we set up and configured a website management interface^[27]^.For the robot application, we implemented the ant nest detection system using ROS (Fig. 6 shows the ant nest detection flow chart). We modified the target tracking module to enable CyberDog to track ant nests, and we set autonomous threshold values based on different backgrounds’ characteristics. Navigation and target confirmation were achieved using Cyclone DDS.

## 3 Result

### 3.1 YOLOX model can recognize different ant nest shapes

The database images were selected randomly according to an 8:2 ratio and used to train the deep neural network while calibrating its parameters. Performance indicators, represented by confidence scores indicating the probability of the model recognizing nests within a picture, were used to evaluate the trained models’ performance in recognizing RIFA nests (Fig. 1a). Results show that different types of RIFA nests were successfully detected and identified. These parameters were found to be significantly higher than the negative controls (Fig. 1b and 1c) after verifying data training. The optimized network model was then tested with 52 RIFA nest images, resulting in a precision rate of 0.95, with only one unrecognizable nest, leading to missing rates of 1.92%. The mean Average Precision (mAP) at a 0.5 Intersection Over Union (IOU) threshold value was 93.44%. Inference time per validation image was 20.16 milliseconds (ms), equivalent to about 50 frames per second, which is a suitable computational speed for CyberDog’s real-time operation.

**Fig. 1.**
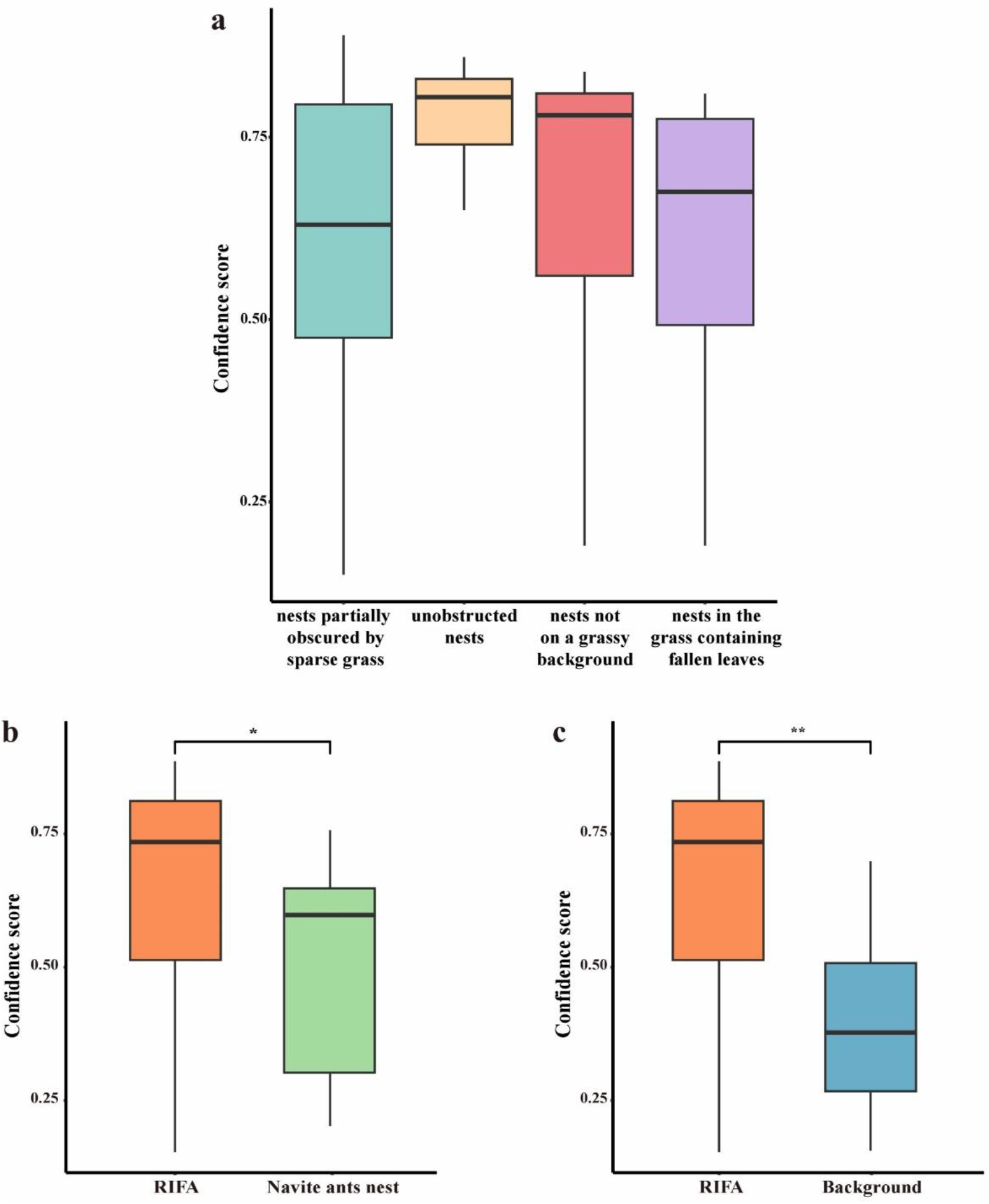
YOLOX recognition efficiency. (a) YOLOX model recognition performance for images of red imported fire ant nests captured under different natural conditions. We evaluated the detection precision on four categories: nests partially obscured by sparse grass (n=43), unobstructed nests (n=16), nests not on grassy background (n=13), and nests on grass land containing fallen leaves (n=10). The results show that our model can effectively detect fire ant nests across various natural environments. (b) The distribution of recognized confidence scores for different ant species. (c) The confidence score distribution comparison between RIFA (red imported fire ants) and background images.

The AI model was trained to distinguish RIFA nest mounds from other surrounding terrain features in the image. Different soil features, surrounding vegetation, and seasonal changes can affect the detection of nests (Fig. 1a). To test the detection efficiency under multiple factors, images of four nest scenarios were gathered and analyzed: partially obscured by sparse grass (n = 43), unobstructed nests (n = 16), nests not on a grassy background (n = 13), and nests in the grass containing fallen leaves (n = 10). The model successfully detected all mounds within the images (Fig. 2 shows some results of RIFA nest detection on test datasets from various locations) with a highest recognition confidence score of 0.5 when dealing with unobstructed nests. Identification results for nests partially obscured by sparse grass, or not in a grassy background, or amidst fallen leaves were also tallied (Fig. 1a). For accurate detection, the model needs to be adjusted using appropriate confidence score cutoff intervals for different nest scenarios.

**Fig. 2.**
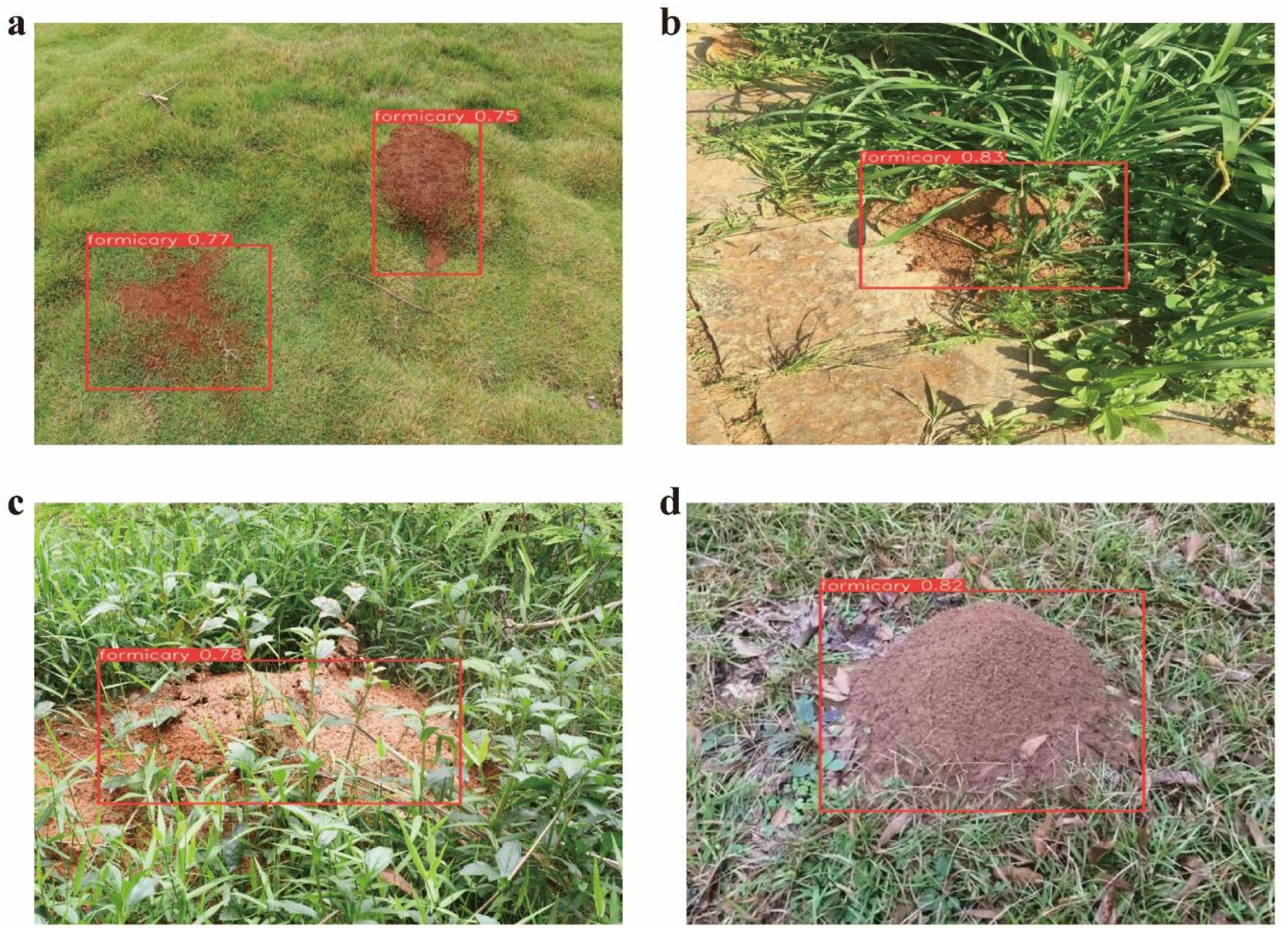
Results of RIFA nest detection on test datasets from various locations. (a) Recognition results for a RIFA nest located within a grassy field. (b) Recognition results for a RIFA nest situated next to a road. (c) Recognition results for a RIFA nest positioned against a high grassland background. (d) Recognition results for a RIFA nest located in a grassland with fallen leaves.

### 3.2 Detection of RIFA nests in the field by the CyberDog installed with the trained model

To evaluate the performance of the YOLOX model for real-time RIFA nest detection, we integrated the trained recognition model into a Cyberdog robot from Xiaomi Co., China. The robot’s built-in identification camera provides real-time footage that is transmitted to the computing platform for nest identification. The Cyberdog, equipped with the YOLOX image recognition algorithm, was used to identify fire ant nests amidst various terrains and vegetation settings (Fig. 3).When the Cyberdog detects a bounding box containing a potential RIFA nest within a certain threshold confidence score, the software initiates multiple measurements of the nest area from two distances(Fig. 4a) - first at 1.2 meters (around twice the typical radius of the mound) from different angles (Fig. 4b and 4c), followed by a closer distance of 0.6 meters for a second round of measurements (Fig. 4d). An active RIFA nest is only confirmed when the second round of measurements shows significantly higher values than the first.

**Fig. 3.**
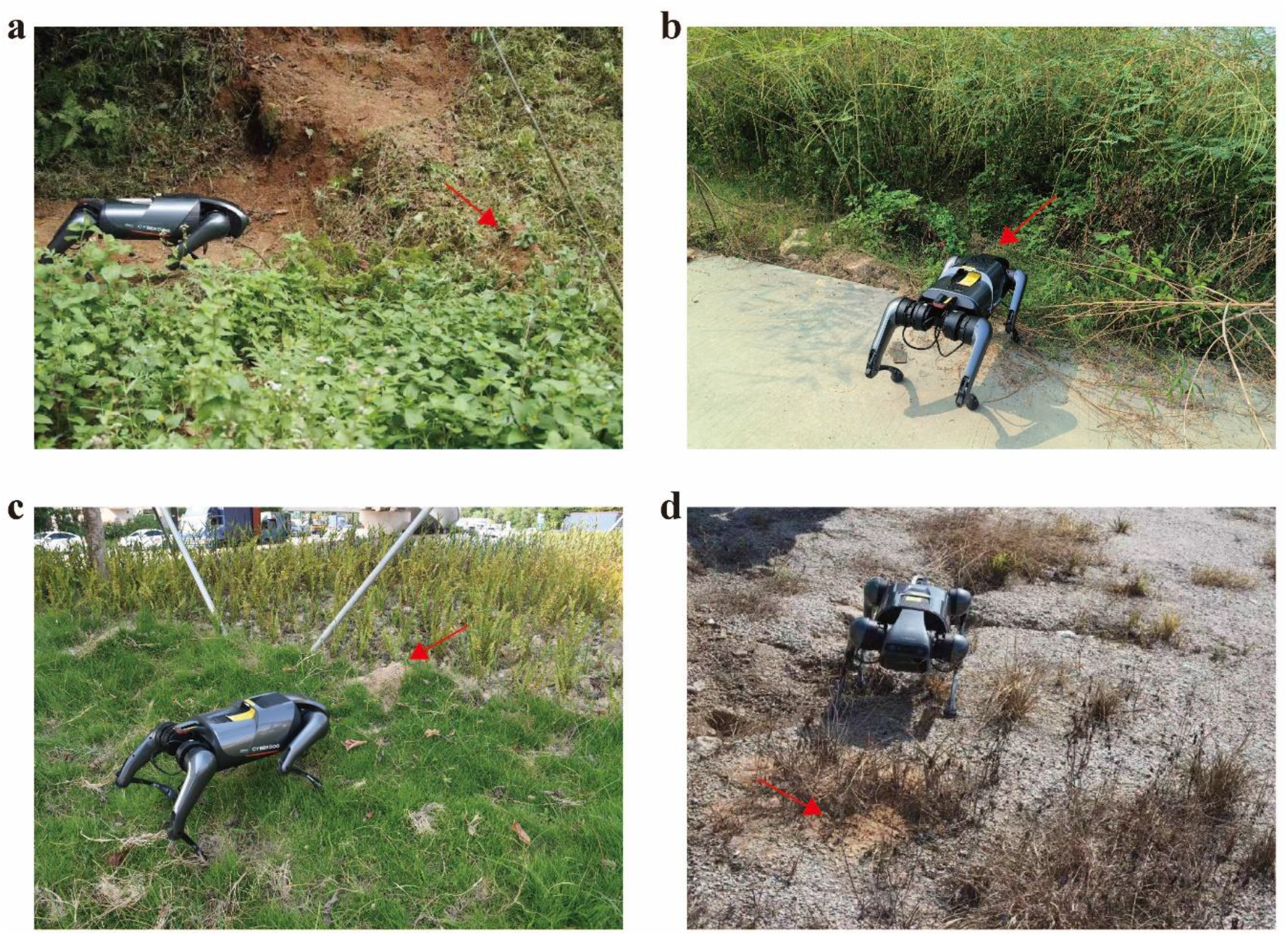
Application of YOLOX image recognition algorithm on the CyberDog for RIFA nest detection in various terrains and vegetation settings. (a) The nest is located on a mountain road slope, surrounded by taller plants. (b) The nest is positioned on a flat concrete road, surrounded by weeds exceeding the CyberDog’s height. (c) The nest is situated on a grassy slope with blades lower than the CyberDog. (d) The nest is located on gravel sand with less vegetation in its proximity.

**Fig. 4.**
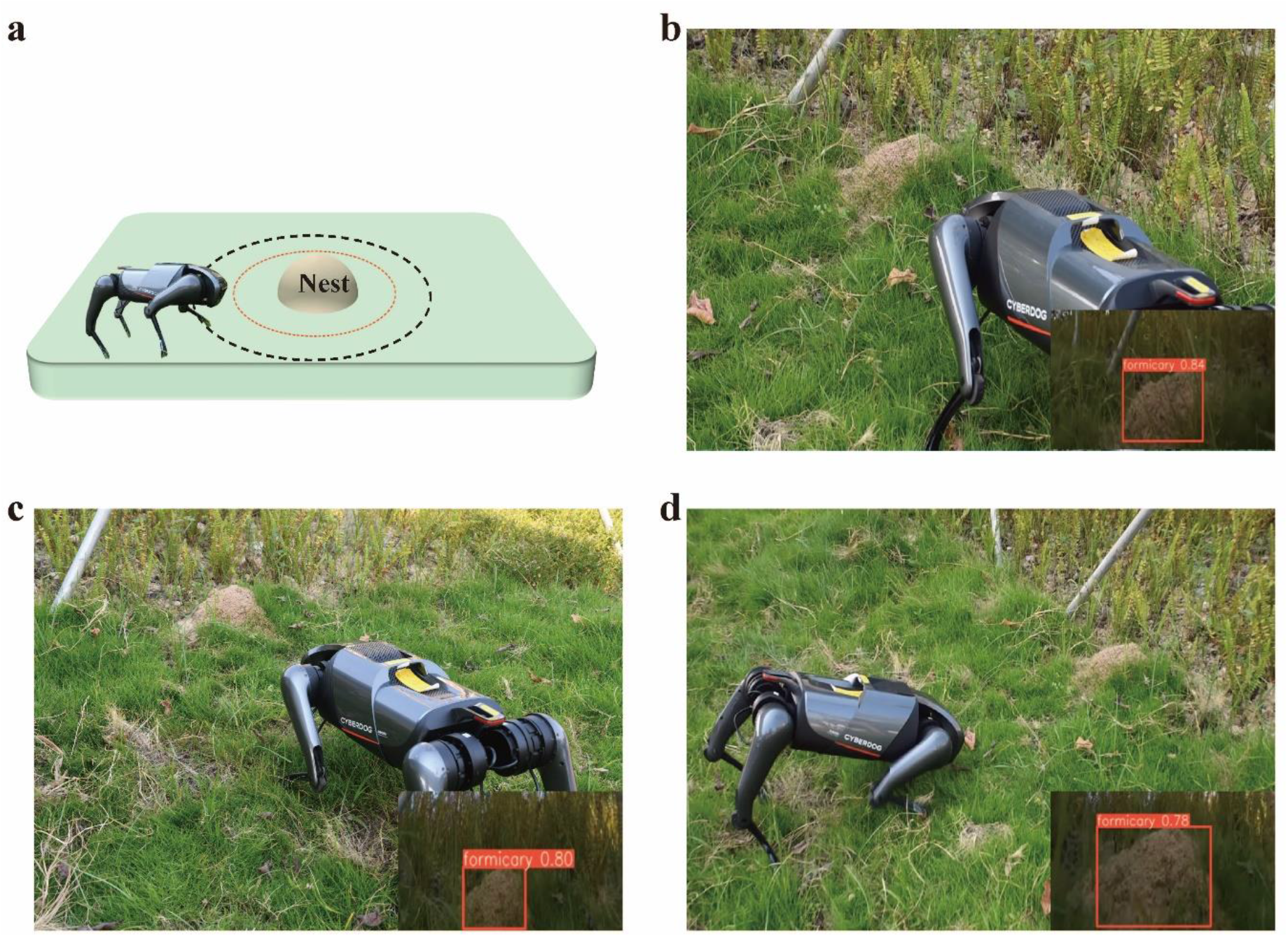
Operational procedure of the CyberDog in detecting a potential RIFA nest. (a) Typical illustration of the CyberDog encountering a RIFA nest. The CyberDog takes multiple measurements from different angles at an initial distance of 1.2 meters (around twice the typical radius of the mound). Then, it moves closer to the target and performs a second round of measurement at a distance of 0.6 meters. If the second round’s measurements are significantly higher than the first, the CyberDog will use its front leg to disturb the nest and check for the presence of ants. (b) Snapshots illustrating the programmed response during target detection at a distance of 1.2 meters. (c) The CyberDog takes multiple measurements to calculate a series of confidence scores. (d) The CyberDog moves closer to the target nest for the second round of measurements to calculate another set of confidence scores. The coordinates of the nest plenum’s location are recorded.

Furthermore, the Cyberdog has been programmed to disturb the nest using its front leg and register any issuing ants’ defensive behavior, such as raised gasters, protruding stingers, and opened mandibles. Highly responsive social insects, fire ants are known to aggressively defend their nests against disturbances. Whenever a mound is disturbed, active fire ants respond by rushing out of any openings in the mound and displaying aggressive behavior. This behavior can be utilized to distinguish active mounds from abandoned ones. Disturbing the nest with the Cyberdog’s front leg significantly enhances recognition accuracy and avoids mixing up with non-fire ant or termite-occupied nests.

### 3.3 Compared detection efficiency of RIFA nests between CyberDog and humans

The robotic dog was developed to either assist or entirely replace humans in detecting RIFA nests with high efficiency. We compared the performance of the CyberDog, programmed for this task, to a typical human inspection. Three people received one hour of training on RIFA biology and identification (similar to the pest management training available in China) before being asked to locate hidden RIFA nests within an open area measuring 300 square meters. We recorded the time spent by both groups patrolling the inspection area, their number of nests successfully found or missed, and the occurrence of misidentifications for all replications. The results indicated that the CyberDog and humans could both complete the task in under ten minutes, with humans being marginally faster, but not significantly so (Fig. 5a, p > 0.05). However, the CyberDog detected three times more RIFA nests than the just-trained humans (Fig. 5b, p < 0.01). In addition, both the missing rate and false detection rate of the CyberDog were significantly lower than those of the humans (Fig. 5c and 5d, both p < 0.01). Thus, our findings suggest that CyberDog equipped with trained recognition model could surpass human detection of fire ants in field surveys.

**Fig. 5.**
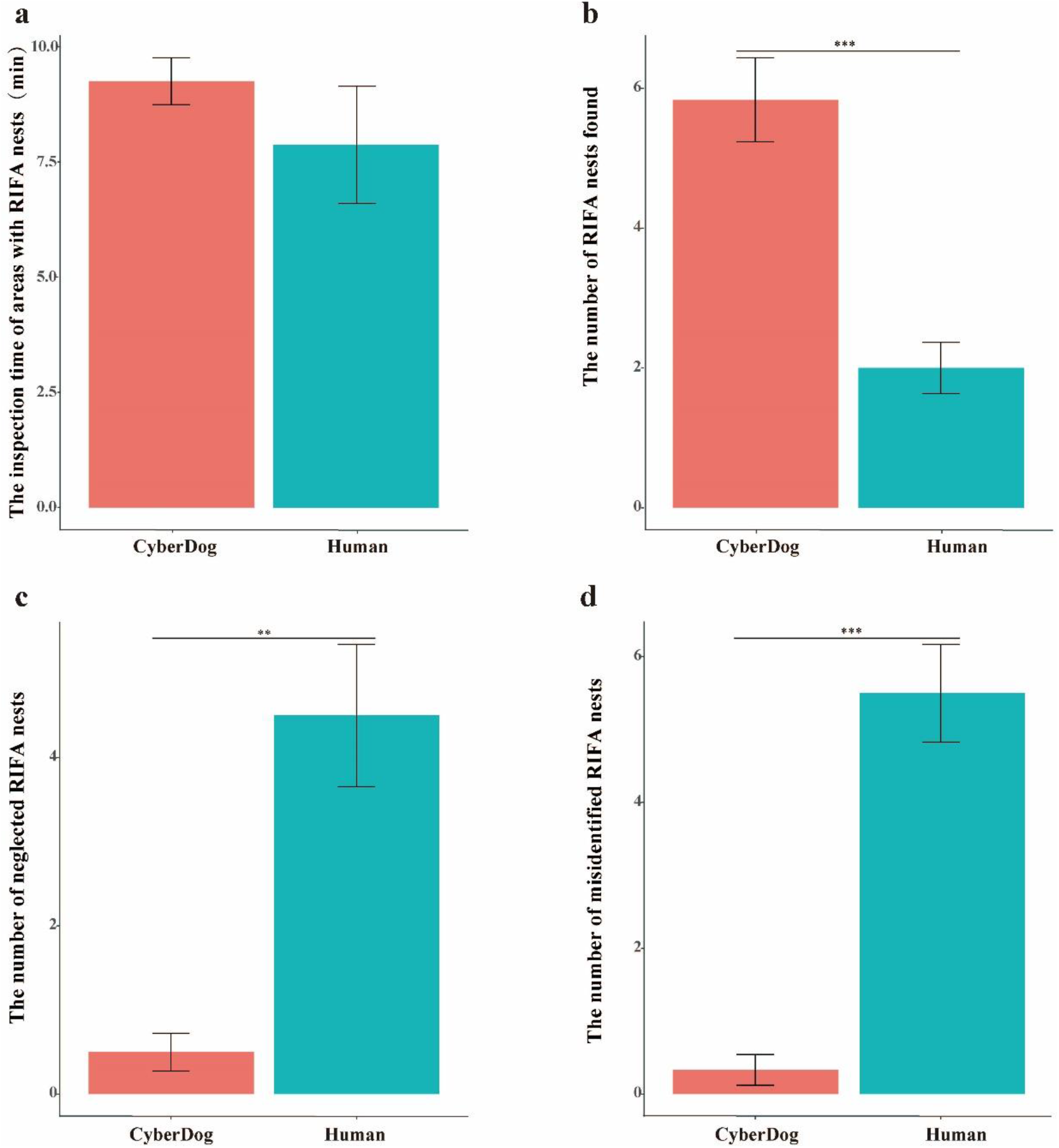
The recognition efficiency of RIFA nests between CyberDog and humans in a test area with known RIFA nest locations was determined by comparing. (a) the amount of time taken to patrol the area, (b) the number of RIFA nests found, (c) the number of RIFA nests missed, and (d) the number of objects misidentified as RIFA nests. The results of the comparison indicated significantly different results between CyberDog and humans, with the P value being less than 0.001 for each of the four measurements.

**Fig. 6.**
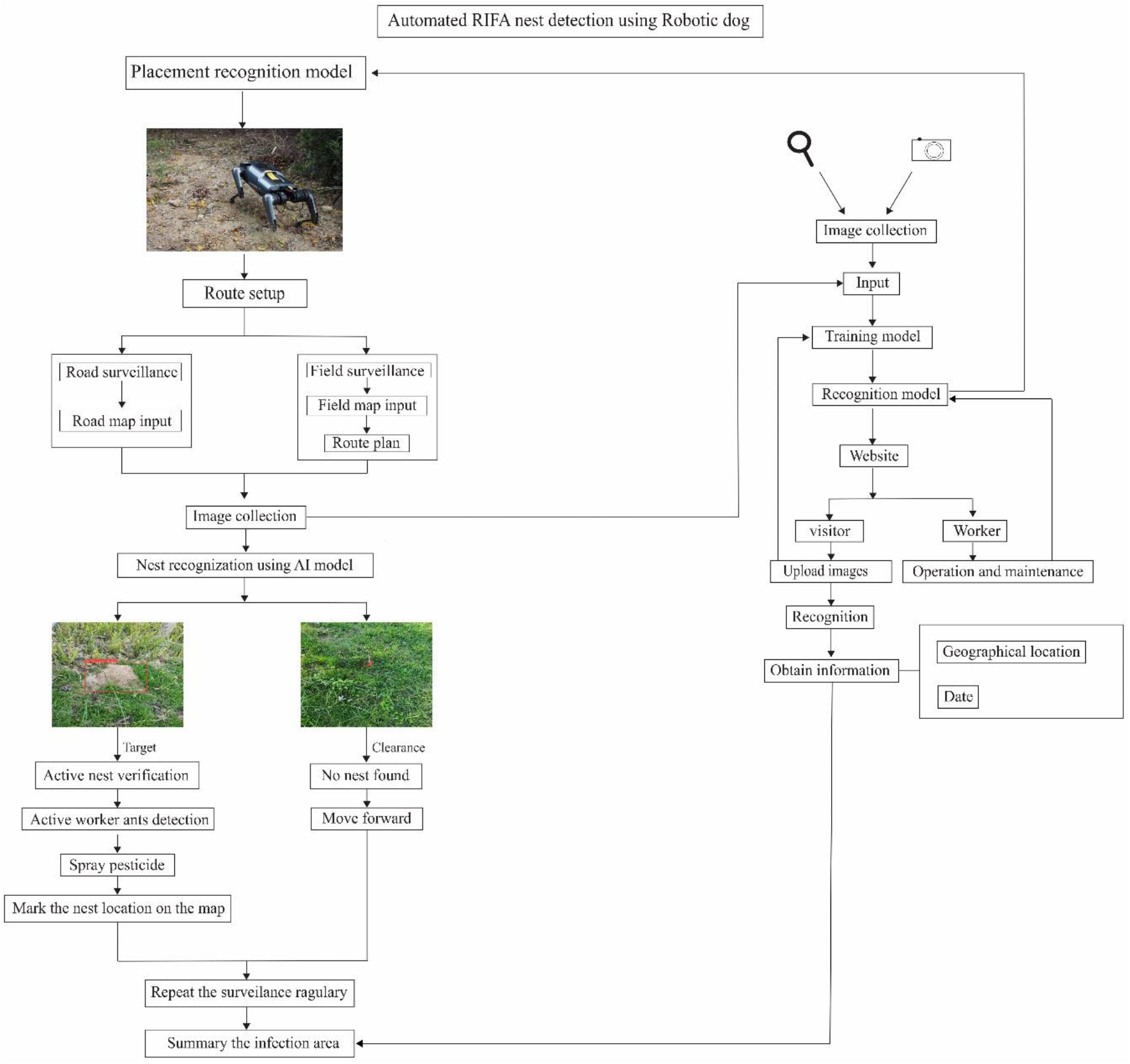
A typical flow chart depicting the RIFA nest detection process using the robot dog.

## 4 Discussion

To enable effective interventions and evaluate the success of suppression efforts, integrated pest management requires periodic monitoring of pest species populations. Prevention is usually less expensive than remediation, making measures that preclude the spread of invasive pests from transportation centers and new infestation sources desirable. Early detection of RIFA infestations can be achieved by surveillance of visible nests in open areas. Various approaches have been evaluated for early detection of RIFA nests over the years. Our method provides a reliable solution for RIFA identification and prevention (Fig. 6). Anyone can upload images to our website for identification at any time, which helps train the model, eventually forming a virtuous cycle. The information collected from these images can be summarized and used for further monitoring and control of the spread of RIFA epidemic areas.

The introduction of image recognition could help to identify RIFA nests towards integrated pest management, thus reducing the amount and costs of pesticide applications, which can greatly cut chemical contamination and damage to local ecology. Simultaneously, the publication of training and testing data sets can make it possible to accelerate the development of other training models implemented with robots using open source computing platforms to help with eliminating pests. The development of robotic dogs as pest management technologies has experienced considerable advancements in the past few decades, providing more efficient and accurate solutions for fire ant pest management. With the help of our open source trained model and datasets, these robotic dogs can now be used in various inspection scenarios, including delivering pesticides to help with fire ant elimination and damage prevention from RIFAs. Moreover, preventive application of some anti-adherent to robotic dog legs may prevent aggressive RIFAs from intruding and damaging internal circuits. Although the current price of the robot dog is relatively expensive, at the same time in order to facilitate action, the power consumption is large, need to be replenished at any time. But with the development of technology and the popularization of robot dogs, these problems will be solved. Robot dogs will be more widely used in the future. In the past decade, the widespread use of unmanned aerial vehicles (UAVs) integrated with sensors and multiple image capture devices have become a new paradigm in remotely sensed imagery. UAV-related technologies brought exciting advancements in the domain of autonomous real-time surveillance, with ever-increasing capabilities in detection, tracking and classification of visualized objects. For example, some pattern recognition algorithms^[28]^ could detect RIFA nests from UAV-obtained imagery. However, it proved difficult to achieve high precision in UAV-borne field inspections that could yield an interactive map of the infested areas for the public. At the same time, China is currently very strict about the management of drones. Civilian drones need to be registered, licensed to fly, and comply with restrictions on flying areas and altitudes. Most importantly, drone technology suffer a high false discovery rate in detecting RIFA nests. Although the different technologies have been integrated with promising results, further extensive field tests are required to consolidate a system with flexible, yet tuned, accuracy and detection limits. All in all, the platform integrating robotics, GIS and pattern recognition technologies is shown to provide an effective, fast and easy tool for fire ant monitoring, particularly in urban areas. Furthermore, the images applied by the robot platform can be made accessible online as a dataset illustrating features diversity range among RIFA nests. Such dataset in itself can be an invaluable resource for better understanding RIFA nesting behavior.

## 5 Conclusions

A YOLOX algorithm-aligned robotic dog-shaped automaton in combination with AI image recognition was tested for the effective detection of RIFA nests in an open field. The robot demonstrated accurate target detection through various terrains, suggesting that multiple such robots could be employed to patrol a designated area and identify fire ant mounds for more efficient pesticide application. As potential developments, these robots could be equipped with toxic baits or in-situ fire ant elimination assets to eradicate RIFA nests from public parks or other smaller areas. Further experiments are necessary to assess the full potential of these new technologies to stop the RIFA invasion in China.

## Author Contributions

Conceptualization, Z.Y. and X.S.; methodology, X.S. and G.S.; software, X.S. and G.S.; validation, Y.H.; formal analysis, X.S., G.S. and Y.L.; investigation, Z.Y., E.G.P.F. and H.Q.; resources, E.G.P. F. and H.Q.; data curation, Z.X. and H.S.; writing-original draft preparation, X.S. and Z.Y.; writing, review and editing, X.S., G.S., W.D. and Z.Y.; visualization, X.S.; supervision, Z.Y.; project administration, Z.Y.; funding acquisition, Z.Y. All authors have read and agreed to the published version of the manuscript.

## Data availability statement

Data is available upon request.

## Acknowledgement

We gratefully acknowledge the technical assistance provided by the IT laboratory of Lan Zhou University’s Computational Engineering and Technology department.

